# Estradiol enhances ethanol stimulation of ventral tegmental area dopamine neuron firing without limiting ethanol inhibition onto those neurons

**DOI:** 10.1101/2020.12.22.423934

**Authors:** Philip S. Lambeth, Amy W. Lasek, Regina A. Mangieri

**Author notes:** corresponding author (RAM).

## Abstract

Females can progress to alcohol and other substance use disorders more quickly than males. The ovarian hormone 17β-estradiol (E2) contributes to sex differences observed in drug use and abuse and may be a principal driver of these differences. However, it is not entirely clear how E2 acts to affect processing of ethanol reward, and several brain regions and mechanisms are implicated. We sought to clarify the role of E2 in modulating the response of ventral tegmental area dopamine neurons to ethanol. To this end, we recorded spontaneous action potentials and inhibitory post synaptic currents from dopaminergic neurons in acute horizontal brain slices from ovariectomized (OVX) dopamine neuron reporter mice (*Pitx3*-eGFP) treated with either vehicle (VEH) or E2. On the basis of prior work, we hypothesized that E2 administration would cause dopamine cells from OVX+E2 animals to show a more substantial ethanol-induced increase in firing rate compared to control animals. Our data confirmed that ethanol stimulation of the firing rate of dopamine neurons from OVX+E2 mice was greater than that of OVX+VEH animals. Further, we hypothesized that the firing rate increase would be accompanied by a concomitant decrease in ethanol stimulated inhibition onto those same neurons. We found that although ethanol caused the expected increase in GABA_A_ receptor-mediated synaptic inhibition in both groups, there was no difference in this response between OVX+E2 and OVX+VEH animals. Our findings lend additional support for the ability of E2 to enhance ventral tegmental area dopamine neuron responses to ethanol and suggest that this effect is not mediated by an E2-elicited suppression of synaptic inhibition.

## Introduction

Female animals respond differently to drugs than their male counterparts [1,2], and understanding the causes of sex differences are critical for advancing our understanding and developing better therapeutics and treatments for substance use disorders. Recently, more attention has been paid to both well-documented and newly-observed evidence of sex differences in animal behavior and physiology, particularly in relation to drug and alcohol consumption. One reason for the growth of interest in this area of alcohol research is that, although women have historically consumed less alcohol than males, this gap has shrunk substantially over the last several decades, and it is now apparent that females are at least equally at risk for alcohol use disorder (AUD) as males [3–5]. For example, Slade et al. [6] found that males born in the early 1900s were 2.2 times more likely to drink and 3.6 times more likely to experience alcohol related harms than their female counterparts. However, males born in the late 1900s were 1.1 times more likely to drink and 1.3 times more likely to experience alcohol-related harms. Moreover, there is evidence that women can escalate to compulsive drinking more quickly than men, and may be more susceptible to alcohol-related cues and relapse [3,7]. Indeed, it well-established that female rodents consume larger amounts of alcohol than males in a variety of drinking protocols, supporting the idea that social, not biological, causes were responsible for the lower rates of human female AUD in the past. Although the same brain regions and neuroanatomical circuits are likely involved in AUD in both sexes, females have been shown to be responsive to different pharmacological manipulations than males [8–10] indicating the engagement of different neurotransmitter and/or neuromodulatory systems.

One brain circuit substantially implicated in the control of ethanol drinking behavior and the action of ethanol overall is the mesolimbic dopamine system [4,11,12]. The proximal mechanism by which ethanol stimulates dopamine release in a major projection target of the ventral tegmental area (VTA) – the nucleus accumbens – appears to be excitation of dopamine neuron action potential firing activity [13]. Within the VTA, there are thought to be direct actions of ethanol on the dopamine neurons, as well as actions on the presynaptic axon terminals of neuronal inputs from various sources [14]. Deletion of dopamine receptors inhibits ethanol drinking behavior [15] while direct manipulations of VTA dopamine neuron and VTA *gamma*-aminobutyric acid (GABA) neuron activity can modulate drinking behavior [16]. These findings indicate a causal relationship between dopamine transmission in the mesolimbic system and ethanol drinking behavior, and place dopamine neurons as an essential component of the mesolimbic system’s response to ethanol. Thus, sex differences in the response of the mesolimbic dopamine system to ethanol may explain, at least in part, sex differences in ethanol drinking-related behaviors.

Gonadal hormones are a major target of interest for exploring physiological mechanisms underlying sex differences in behavior [9,17]. Females produce ovarian-derived estrogens: estrone, estriol, and estradiol (also known as E2) – which is the most prevalent and well-studied of the endogenous estrogens. Recent studies have examined the role of E2 in alcohol-related behaviors, demonstrating that high levels of E2 increased binge-like ethanol drinking and the rewarding properties of ethanol in female mice [10,18,19]. In one report, [20] found that when mice were in diestrus (with rising levels of E2), the firing activity of VTA dopamine neurons responded more substantially to ethanol than neurons from mice who were in estrus (with low E2). Similarly, when E2 was administered to mice with their ovaries removed (OVX), the dopamine neuron firing response to ethanol was potentiated.

E2 acts on estrogen receptors (ERα, ERβ, and G protein-coupled estrogen receptor), which are expressed in numerous regions throughout the rodent brain, including extensively in the mesolimbic reward pathways [21,22]. Prior work has shown that activation of both ERα and ERβ may be important for the rewarding properties of ethanol and that while activation of each separately was not sufficient, activation of both together can lead to increased ethanol conditioned place preference [10]. In a follow up study, Vandegrift et al. [23] found that selective activation of ERα (but not ERβ) potentiated the stimulation of dopamine neuron firing by ethanol and that knockout of ERα from dopaminergic neurons eliminated the ability of E2 to enhance ethanol stimulation of dopamine neuron firing, suggesting that ERα mediates E2’s ability to promote ethanol stimulation of the mesolimbic system. Finally, Vandegrift et al. also determined that reducing levels of ERα in the VTA of female mice resulted in decreased binge-like ethanol intake. Thus, this body of work indicates that E2-ERα signaling in the VTA promotes ethanol drinking by females, likely by enhancing ethanol stimulation of mesolimbic dopamine signaling.

ERα has classically been considered a nuclear hormone receptor [24,25], with its activation leading to engagement of transcriptional machinery and alterations in gene expression. However, another type of estrogen receptor activity has been described as well; Arnal and colleagues [26] made the distinction between transcriptional activities and rapid, non-genomic, membrane-initiated steroid signals (“MISS”), and we adopt this terminology here. Through associations with other cell membrane receptors, activation of ERα can lead to a variety of downstream effects via second messenger systems. For example, in the hippocampus, ERα and group I metabotropic glutamate receptors (mGluRs) form complexes at the cell membrane via caveolin proteins, after which signaling partners of mGluR1 [27–29], such as the inositol triphosphate (IP_3_) receptor, can be activated by E2 stimulation of ERα [30]. This cascade leads to the postsynaptic production of an endocannabinoid which then acts as a retrograde messenger at cannabinoid CB_1_ receptors on presynaptic GABAergic terminals to suppress GABA release [30].

We hypothesized that a similar MISS-type activity (E2-suppression of inhibition) could account for the observed effect of E2 on ethanol stimulation of dopamine neuron firing. In vitro application of ethanol has been shown to increase the firing rate of some populations of dopamine neurons, but, paradoxically, also to increase inhibitory currents onto those neurons [31,32]. Moreover, it was demonstrated that GABA_A_ receptor antagonism potentiated ethanol stimulation of firing [33] or attenuated ethanol inhibition of firing [32], indicating that a concurrent increase in GABA release elicited by ethanol can limit the excitation of, or inhibit, dopamine neuron firing activity. Therefore, we speculated that E2 might facilitate ethanol stimulation of firing via MISS activity that suppresses GABA transmission onto dopamine neurons.

In summary, prior work indicates that E2 can act to promote drinking behavior in females via facilitation of dopamine neuron responses to ethanol. In general, it is known that several processes converge onto dopamine neurons and combine to mediate ethanol’s effect on firing rate, but the specific mechanism(s) by which E2 acts to facilitate ethanol excitation of firing are not known. Here, we focus on suspected MISS elicited by E2, which we hypothesized would suppress GABA signaling and thereby disinhibit dopamine neurons. We used ovariectomy (OVX) followed by injections of E2 or vehicle (VEH), rather than using freely cycling animals, in order to directly probe the role of estradiol. Firing rate and synaptic GABA responses to ethanol were recorded from VTA dopamine neurons using ex vivo brain slice electrophysiology. Although we replicated the prior finding of E2-potentiatation of ethanol stimulation of VTA dopamine neuron firing activity, we did not observe any effects of E2 on GABA transmission in these neurons.

## Materials and methods

### Animals

All procedures were carried out in accordance with the regulations put forth by Association for Assessment and Accreditation of Laboratory Animals (AALAC) and were approved by The University of Texas at Austin Institutional Animal Care and Use Committee (IACUC). All mice used for these experiments were female, hemizygous knock-in transgenic mice in which the expression of a green fluorophore (enhanced green fluorescent protein; eGFP) was driven by the transcription factor *Pitx3*, which, in the midbrain, selectively marks dopamine neurons (*Pitx3*-eGFP) [34–36]. *Pitx3*-eGFP mice are reported to have very high specificity in expression (∼95% of GFP^+^ cells in the VTA are TH^+^) and only appear to exhibit negative consequences of transgene insertion in homozygotes [34,37,38]. Original stock of two double hemizygous males was obtained from Dr. Chrissa Kioussi (Oregon State University) and this line was maintained by our lab at The University of Texas at Austin by mating double hemizygous transgenic sires with wild type C57BL/6J females. The animals used in this study were the result of at least six generations of backcrossing of the transgenic line with C57BL/6J mice. Mice were group housed from weaning until surgery, and maintained throughout their life on Sani-Chips wood bedding (PJ Murphy). Water and standard chow (LabDiet® 5LL2 Prolab RMH1800) were available *ad libitum* throughout the experiment. After surgery, mice were maintained in single housing adjacent to their former cage mates until the recording day.

From birth until ovariectomy surgery, animals were kept on a 12 h light / 12 h dark cycle with lights on at 7:00 AM. One-three days after surgery, animals were moved to a satellite housing room within our lab. The light cycle was again 12 h light / 12 h dark, but with lights on at 9:30 PM to allow for experiments during the dark phase.

### Ovariectomy

Ovaries were removed from mice that were 8-11 weeks old. After OVX mice recovered from surgery for 1-4 weeks. Surgeries were performed similarly to those described previously [18,20,23] with some procedural modifications detailed below. Briefly, the animals were anesthetized with 3% isoflurane and anesthesia was maintained at 2-2.25% throughout the procedure, which lasted between 45-75 minutes. The animal was placed ventral side down and after the back was shaved and sterilized with Povidone-iodine a ∼12.7 mm vertical incision was made along the dorsal midline centering on the kidneys. This incision was pulled first to one side then the other, to allow access and make a small incision in the muscle wall surround the uterine horn. On each side, the horn was carefully pulled out and fat pads were dissected away before ovary was removed (via clamping and cutting) and uterine horn reinserted into the body. One to two dissolvable sutures were placed internally to close the muscle wall on each side, as needed, and 4-5 nylon external sutures were used to close the wound. Care was taken to induce as little damage to the muscle and fat pads as possible and to minimize pain and suffering throughout. Meloxicam® (0.5mg/kg) (Animal Health International) or Rimadyl® (0.5 mg/kg) were used for pain relief, and Neosporin® as needed for wound maintenance. The animals were monitored closely for post-operative surgical complications.

### E2 Injections

After full recovery from the surgery and adjustment to the new light cycle (> 9 days), mice were treated for three days with 0.05 mL solution containing 10% ethanol in sesame oil (Sigma) vehicle (VEH) or estradiol benzoate (E2, Sigma) in VEH subcutaneously. For the first two days of injections, the animals received 0.2 μg EB or VEH, which has been shown to lead to serum E2 levels approximating those in proestrus at 4 hours after injection [20]. On the third day, 60-75 minutes before the brain slice preparation, the animals were injected with 1.0 μg EB. For 90% of all injections, the experimenter was blinded to the solution administered.

### Verification of OVX

In a subset of the animals, vaginal swabs were taken over the course of 4 or 5 days beginning after the acute surgical recovery period. The swabs were smeared onto individual slides and stained with Wright-Giemsa stain to visualize cellular content. Slides were reviewed by 2-3 observers; if cell type composition was determined to be variable over time, the animal was determined as likely still cycling and its data was not used in the statistical analysis [39,40].

In another subset of animals, the uterus was exposed at the time of brain slice preparation, photographed, and, in some cases, removed and weighed. After a successful OVX, the uterus atrophies substantially [41], although to a lesser degree in our study, given the relatively short length of time between the OVX and electrophysiology experiment. We rejected from analysis animals whose uterus did not show signs of atrophy (in these experiments, average weight of atrophied = 0.045 g, average weight of non-atrophied/failed OVX = 0.101 g). Overall, 100% of the animals included in the firing rate analysis were verified via swab; for the IPSC experiment, 6 animals were verified by swab only, 5 animals by uterus atrophy only, and 4 animals using both methods.

### Preparation of Brain Slices

Slices were prepared using methods adapted from [42] and [33]. Mice were anesthetized by inhalation of isoflurane (open-drop) and quickly decapitated using a guillotine. The head was immediately submerged into ice cold oxygenated sucrose-based artificial cerebrospinal fluid (aCSF) containing: (in mM) 210 sucrose, 26.2 NaHCO_3_, 1 NaH_2_PO_4_, 2.5 KCl, 10 dextrose, 7 MgSO_4_, and 0.5 CaCl_2,_ bubbled with 95% O_2_/5% CO_2_. One minute later, the brain was rapidly removed and then blocked to prepare for horizontal slicing. Horizontal sections (190 μm thick) were made through the midbrain, collecting 2 to 3 slices containing VTA. They were then transferred to an oxygenated recovery solution at 33°C containing: (in mM) 120 NaCl, 25 NaHCO_3_, 1.25 NaH_2_PO_4_, 2 KCl, 10 dextrose, 2.4 MgSO_4_, and 1.8 CaCl_2_, where they were maintained for at least 30 minutes and up to several hours before being transferred to the recording apparatus.

### Patch Clamp Electrophysiology

Cell-attached current- and voltage-clamp recordings (firing rate) and whole-cell voltage clamp recordings (inhibitory post synaptic currents, IPSCs) were conducted in the lateral VTA, informed by their location in relation to the medial nucleus of the accessory optic tract (MT) [33,43] (see Fig 1). Cells were identified by BX50 microscope mounted on a vibration isolation table. Fluorescence was verified by epifluorescent illumination of eGFP using the MRK200 Modular Imaging system from Siskiyou Corporation (see Fig 1 for representative images). Recordings were made in an aCSF that contained: (in mM) 120 NaCl, 25 NaHCO_3_, 1.25 NaH_2_PO_4_, 2 KCl, 10 dextrose, 1.2 MgSO_4_, and 2.0 CaCl_2_, continuously bubbled with 95% O_2_/5% CO_2_. ACSF was constantly perfused at a rate of ∼2mL/min and maintained at a temperature of 30-33° C via an inline bath heater (Warner Instruments). All drugs were purchased from Sigma except where otherwise noted.

**Fig 1.**
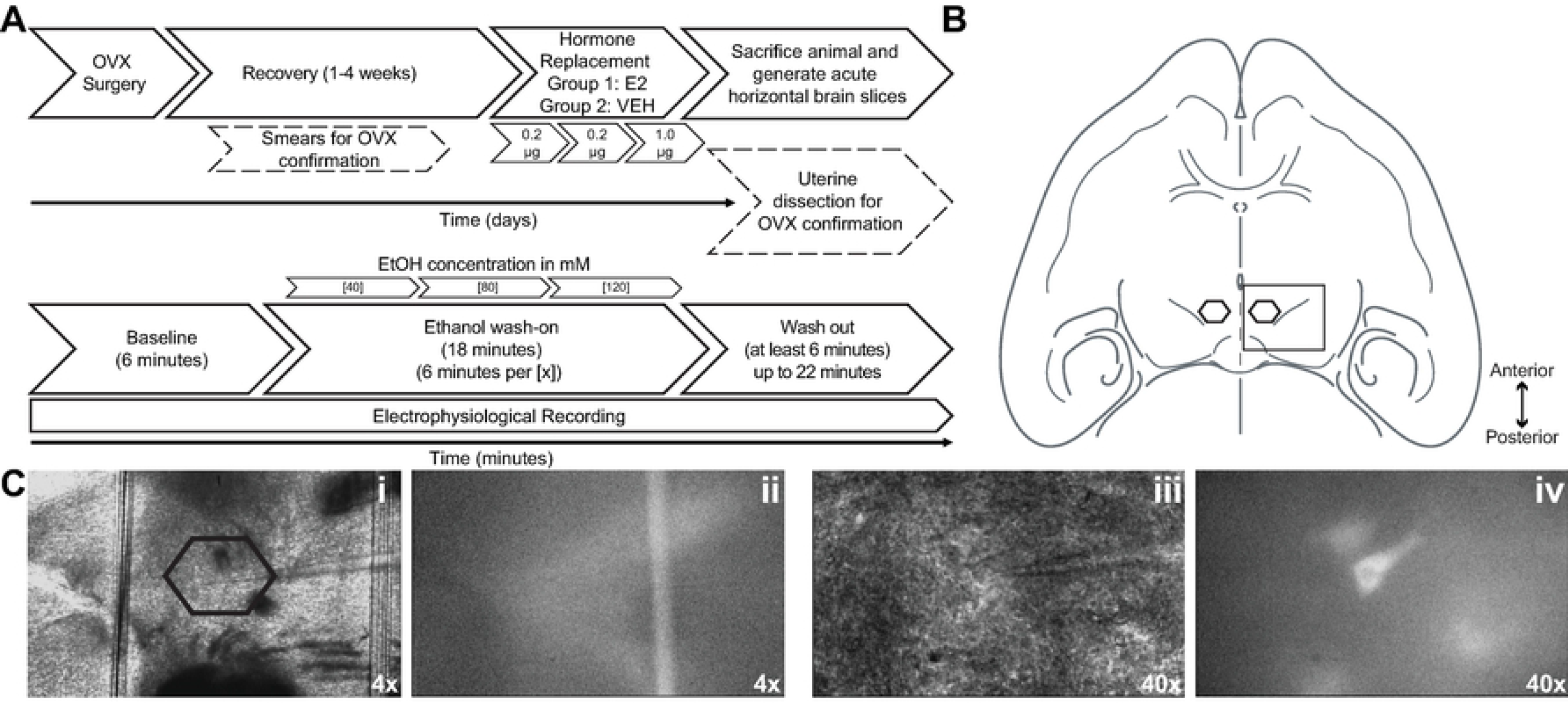
Experimental timeline and methods. (A) Experimental timeline (top), in days, followed by electrophysiology recording timeline (below), in minutes. (B) Cartoon depicting horizontal brain slice containing the VTA; our recordings often also consist of one 190 μm slice dorsal to this and one slice ventral. Hexagonal shapes indicate regions from which cells were selected for recordings. Boxed region corresponds to 4X images shown in panel C. (C) Example images of brain slice and electrode placement during electrophysiology recordings. (i) widefield @ 4x magnification, MT visible near center, with electrode. Hexagonal shape corresponds to same region indicated in panel B. (ii) eGFP fluorescent version of same field as (i). (iii) widefield @ 40x, electrode and cell visible. (iv) eGFP fluorescent version of same field as (iii). MT: medial terminal nucleus of the accessory optic tract.

For whole cell voltage clamp recordings, DNQX (50 μM), APV (20 μM), and CGP52432 (1 μM) were included in the recording solution to block AMPA, NMDA, and GABA_B_ receptors, respectively. Recording electrodes (4” thin-wall glass capillary tubes; 1.5 OD / 1.12 ID – World Precision Instruments) were made using a P-97 Flaming/Brown™ micropipette puller (Sutter Instruments) to yield resistances between 3.5–6 MΩ and were filled with an internal solution consisting of: (in mM) 100 KCl, 50 K-gluconate, 10 HEPES, 0.2 EGTA, 4 Mg^2+^-ATP, Na^+^-GTP, Phosphocreatine disodium salt hydrate, or 145 KCl, 10 HEPES, 5 EGTA, 5 MgCl_2_, 2 Na-GTP, 0.2 Na-ATP (one recording only). For cell attached recordings, electrodes were made using a PC-10 dual-stage glass micropipette puller (Narishige) to yield resistances between 2 and 4 MΩ and were filled with normal recording aCSF.

For confirmation of the nature of the spontaneous IPSCs (sIPSC), two recordings were performed in which picrotoxin (50 μM, Sigma) was added at the end of the wash period. In both analyzed cells, the application of picrotoxin eliminated ≥99% of sIPSCs, indicating that these events are predominantly GABA_A_-R mediated.

### Electrophysiology Data Acquisition

All voltage and current responses were acquired using CV-7B headstage with MultiClamp 700B amplifier, filtered at 1 kHz (firing rate) or 2 kHz (IPSCs), and digitized at 10 kHz via a Digidata 1440A interface board using pClamp 10 (Axon Instruments). All recordings were spontaneous events, whether in cell-attached mode for firing rate experiments, or whole cell for the IPSC experiments. IPSCs were monitored with a −60 mV holding potential. For the firing rate recordings, after a loose seal (5–25 MΩ) was formed, 10 minutes of stable baseline was acquired, at which point the first concentration of ethanol was washed on to the slice. The ethanol-containing solutions were freshly prepared in the recording aCSF using 95% ethanol (Pharmco-Aaper) to final concentrations of 40, 80, and 120 mM. After the baseline period, each concentration was successively applied for 6 minutes, after which a wash period of at least 6 and up to 20 minutes was recorded. The IPSC recordings were performed as described above, except that after forming a tight seal (> 1 GΩ), a current clamp recording of 2 to 3 minutes was acquired (to monitor the firing rate) after which the cell was broken into by application of slight negative pressure. Immediately after break in, I_h_ was measured by administering a 1.5 s hyperpolarizing pulse from −60 mV to −120 mV and measuring the amplitude of the sag current at steady state. Following, the baseline spontaneous IPSC recording began and proceeded as with the firing rate, with the wash period consisting of at least 6 minutes. Spontaneous acquisition was performed in sweeps of 9.375 s repeated every 10 s, with a 0.625 s period for membrane test between sweeps. Once any ethanol had been applied to the slice, no additional cells in that slice were used for recordings. For all whole cell recordings, series resistance was monitored throughout the experiment and we rejected those cells for which the variability was greater than 30%, measuring from the time at which ethanol reached the bath. Events were detected using either the Threshold Search (AP firing) or Template Search (sIPSCs) features in ClampFit (Axon Instruments); sIPSC events smaller than 10 pA in amplitude were excluded from analysis. Search results (event times, and peak amplitudes for sIPSCs) were transferred into Microsoft Excel and examined for quality control by inspecting anomalous interevent intervals to identify and remove spurious events before being processed in MATLAB (Mathworks) to produce average frequency and amplitude per sweep. These were then used to derive the one-minute bin averaged values. The experimenter was blinded to the treatment that the animal received from injections throughout the acquisition and analysis in 90% of cases; there were 3 animals total who comprised part of the final analysis (2 E2 and 1 VEH) whose treatments were known to the experimenter throughout the experimentation and analysis.

### Data presentation and Statistical Analysis

Baseline comparisons were planned via t-test. Data were tested for normality. In some cases, the baseline data were non-normally distributed, or the variances were not equal, and non-parametric tests or Welch’s t-test were used, and are indicated, where appropriate. Group summary data show mean ± SEM for normally distributed data; median and interquartile ranges for non-normally distributed data. Time course data were expressed as percent change from the 6-minute, pre-ethanol application, baseline average; figures show these data plotted in 1-minute time bins. The firing rate time course was analyzed by two-way repeated measures mixed ANOVA, between-factor – Treatment (E2, VEH), within-factor – Concentration/Phase (40, 80, 120, wash), using the 6-minute average for each Concentration/Phase; sIPSC time course analyses were identical, except they did not include the wash period. Sphericity was not assumed and, if shown to be violated as calculated by Mauchly’s test, the Greenhouse-Geisser corrected degrees of freedom and p-values were used and are reported in the text. Sidak’s post-hoc tests comparing treatment groups within each phase were performed in the case of a significant interaction effect. IBM SPSS Statistics 25 and Prism 8.0 were used to perform statistical analyses. A p-value of < 0.05 was considered significant and exact values are reported to three significant digits.

## Results

### E2 enhanced ethanol excitation of VTA dopamine neuron firing

We first verified that we could replicate the effect of E2 described in [20] given several differences in our electrophysiology methodology and the line of mice used. As described in more detail in the Materials and methods section, *Pitx3-eGFP* mice underwent an OVX surgery, followed by a recovery period of 1-4 weeks. For two days prior to the recording day, mice received daily injections of E2 (0.2 µg) or VEH, with a larger dose (1 µg) of E2 administered on the third day (Fig 1A). Beginning 60-75 minutes after the third injection of E2 or VEH, brain slices containing the VTA were prepared for electrophysiological recordings and action potential firing frequency (Hz) of VTA dopamine neurons was determined by recording spontaneous voltage deflections of eGFP^+^ cells in loose-seal configuration (Figs 1B,C and 2A). During the 6-minute baseline period prior to ethanol application, the firing rate of cells from E2-treated mice was not different from VEH-treated (Komolgorov-Smirnov D = 0.2556, n.s.) (Fig 2B). We then measured the response to increasing concentrations of ethanol in the extracellular fluid (Fig 2C). Similarly to Vandegrift et al. [20] we found that ethanol application elicited a greater increase in the firing rate of cells from E2-treated mice (two-way repeated measures mixed ANOVA, main effect of Treatment: F(1,17) = 13.52, p = 0.0019). We also observed that the effect of Concentration (Geisser-Greenhouse corrected F(1.708, 29.03) = 10.50, p = 0.0006) varied with the treatment condition (Treatment x Concentration interaction (F(3,51) = 3.815, p = 0.015). These results indicate that while ethanol overall elicited a concentration-dependent increase in firing frequency, dopamine cells from E2-treated animals were more excited by ethanol.

**Fig 2.**
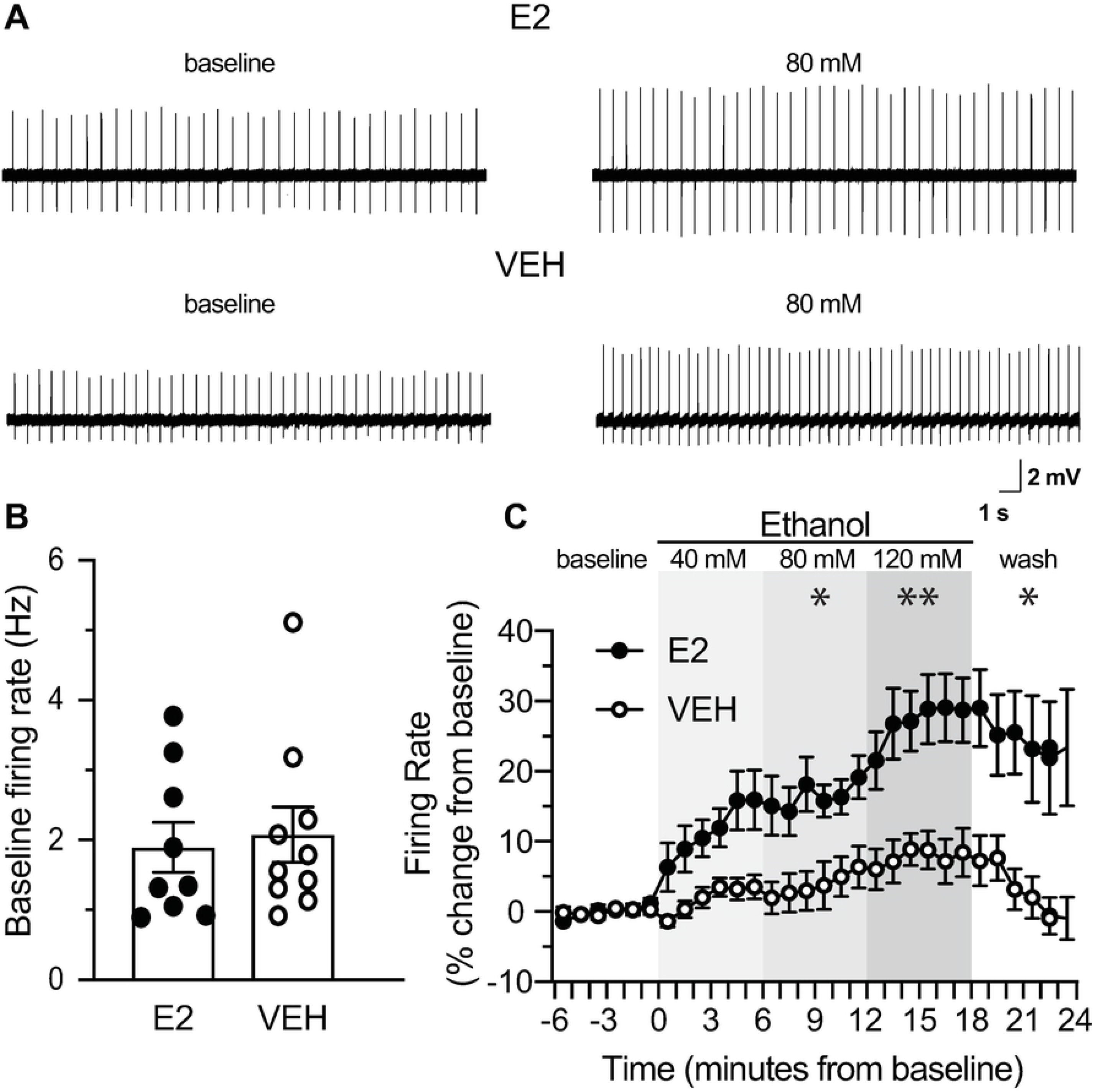
Estradiol treatment potentiates ethanol-stimulated firing in VTA dopamine neurons. (A) Example voltage traces showing 10 s each of baseline and 80 mM ethanol. (B) Baseline firing rates during the 6-minute period prior to bath ethanol application were not different between treatment groups (Welch’s t = 0.6291, n.s.). E2 (estradiol): n = 9 cells from 5 mice, VEH (vehicle): n = 10 cells from 5 mice. Circles show data for individual cells, bars depict group means ± SEM. (C) Firing rate over time displayed as percent change from baseline average in one-minute bins. Ethanol application caused a concentration-dependent increase in firing rate overall (p = 0.0006), with a Treatment x Concentration interaction (p = 0.015). Sidak’s post hoc tests indicated significantly higher firing rate in the E2 group than VEH during 80 mM (p = 0.023), 120 mM (p = 0.009) and wash (p = 0.042) phases, but not during 40 mM (p = 0.072). *, p < 0.05, **, p < 0.01, for within-phase comparisons of E2 versus VEH. E2: n = 9 cells from 5 mice, VEH: n = 10 cells from 5 mice. Circles show group means ± SEM.

### E2 does not affect ethanol to increase sIPSC frequency onto VTA dopamine neurons

Next, we tested our hypothesis that the firing rate differences described above were the result of a different GABA response to ethanol in cells from E2-treated mice compared to VEH-treated. To examine this question, we performed whole-cell recordings in VTA dopamine neurons and measured spontaneous GABA_A_ receptor-mediated IPSCs (Fig 3A). The experimental design was similar to that of the firing rate experiment described above, but with added steps as the whole cell recordings require, including monitoring of cell membrane properties. I_h_ is a current mediated by hyperpolarization-activated cyclic-nucleotide gated channels in dopamine neurons and has been used historically to identify putative dopamine neurons [12,36,44,45]; although we relied on fluorescence here for identification, we did measure this current immediately after rupturing the cell membrane and achieving whole-cell recording configuration. We measured an observable I_h_ in all recorded cells, and E2 treatment did not appear to affect the magnitude of I_h_ (E2: 127.6 ± 21.53 pA, VEH: 129.8 ± 39.45 pA; Kolmogorov-Smirnov D = 0.2095, p = 0.927). When analyzing IPSCs, as with the firing rate, there were no significant differences between neurons from E2- and VEH-treated animals at baseline (Fig 3 B,D,E; frequency (Hz): unpaired t-test: t=0.5362, n.s.; amplitude (pA): Welch’s t = 0.6291, n.s.; drive (amplitude x frequency): Kolmogorov-Smirnov D = 0.3463, n.s.). Furthermore, we did not observe any differences between E2- and VEH-treated groups in the time course for frequency as ethanol was applied (Fig 3C, two-way repeated measures mixed ANOVA, no main effect of Treatment: F(1,19) = 0.855, p = 0.367; no interaction: F(2,38) = 0.273), although there was a clear effect of ethanol to increase IPSC frequency (main effect of Concentration: F(1.3, 23.7) = 24.91, p < 0.0001). Similarly, we observed no differences in amplitude between E2- and VEH-treated groups (Fig 3E; two-way repeated measures mixed ANOVA, no main effect of Treatment: F(1,19) = 0.265, p = 0.613; no interaction: F(2,38) = 0.276, p = 0.761) although there was a small effect of ethanol to increase amplitude over time (main effect of Concentration: F(1.336, 25.39) = 5.936, p = 0.015). Finally, the IPSC drive time course was similar to the IPSC frequency (Fig 3G; two way repeated measures mixed ANOVA, no main effect of Treatment: F(1,19) = 1.215, p = 0.284, no interaction: F(2,38) = 0.637, p = 0.534, main effect of Concentration: F(1.096, 20.81) = 18.51, p = 0.0002). This indicates an increased inhibitory drive in response to ethanol, regardless of in vivo treatment condition, although this effect on GABA signaling appears to be mainly driven via the increased frequency. Therefore, we did not observe evidence that E2 acts to potentiate the dopamine neuron firing rate response to ethanol through modulation of the response of GABA afferents to ethanol.

**Fig 3.**
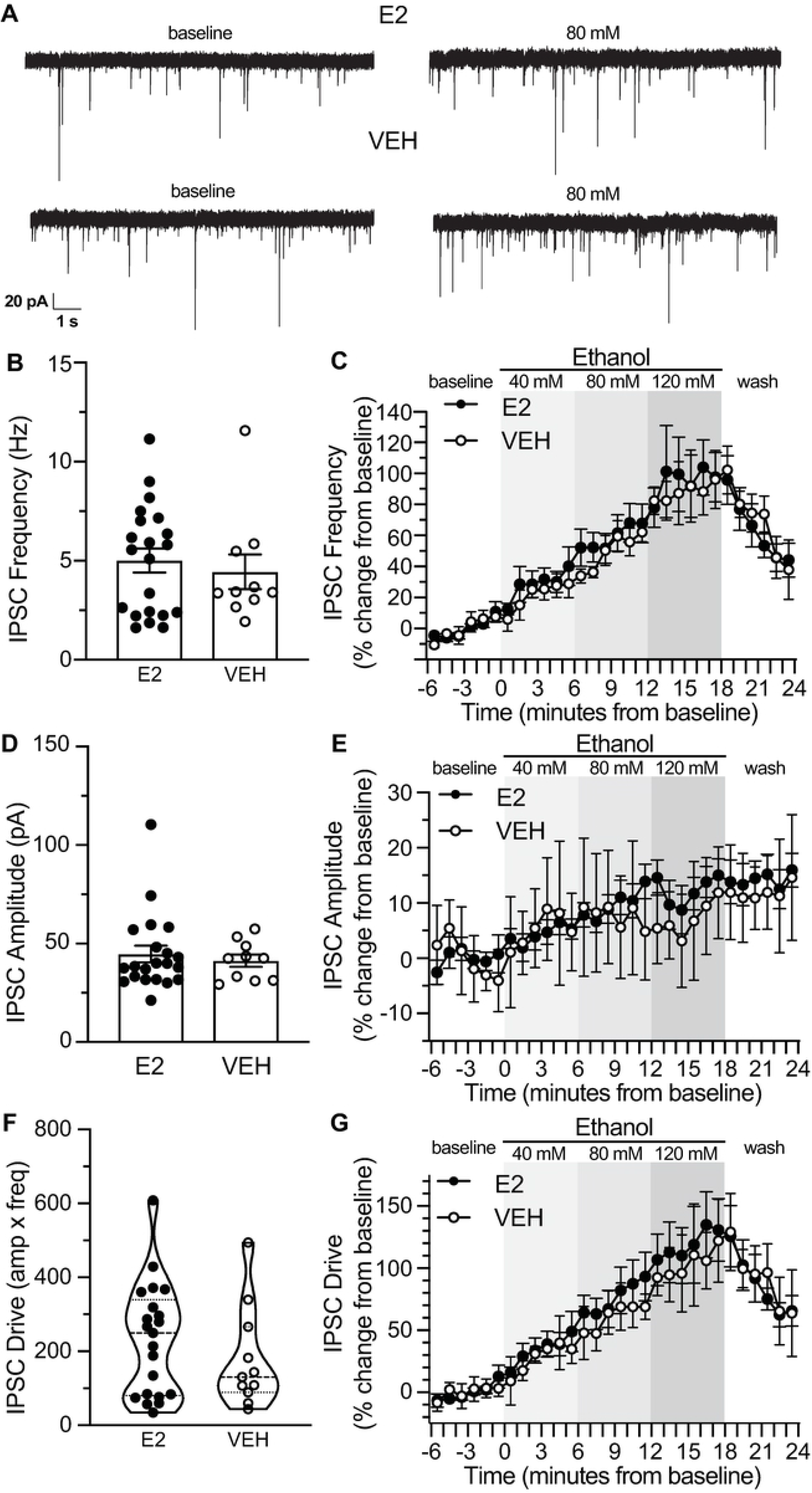
Estradiol treatment did not affect ethanol enhancement of GABA_A_R-mediated transmission ex vivo. (A) Example current traces showing 10 s each of baseline and 80 mM ethanol. (B) Baseline inhibitory postsynaptic current (IPSC) frequency during the 6-minute period prior to bath ethanol application was not different between groups (t = 0.5362, n.s.). (C) IPSC frequency over time displayed as percent change from baseline average in one-minute bins. Ethanol application caused a concentration-dependent increase in IPSC frequency that did not differ between groups (main effect of Concentration, p < 0.0001, no effect of Treatment or Treatment x Concentration interaction). (D) Baseline IPSC amplitude was not different between groups (Welch’s t = 0.6291, n.s.). (E) IPSC amplitude over time, displayed as in (C). Ethanol application caused a concentration-dependent increase in IPSC amplitude that did not differ between groups (main effect of Concentration, p = 0.015, no effect of Treatment or Treatment x Concentration interaction). (F) Baseline IPSC drive was not different between groups (Kolmogorov-Smirnov D = 0.3463, n.s.). (G) IPSC drive over time, displayed as in (C). Ethanol application caused a concentration-dependent increase in IPSC drive (main effect of Concentration, p = 0.0002, no effect of Treatment or Treatment x Concentration interaction). For baseline comparisons (B, D, F), E2: n = 21 cells from 13 mice, VEH: n = 10 cells from 7 mice; circles show data for individual cells, bars depict group means ± SEM and violin plots show group medians and interquartile ranges. For time course analyses (C, E, G), E2: n = 15 cells from 11 mice; VEH: n = 6 cells from 4 mice; circles show group means ± SEM. Not all cells used in the baseline comparisons were included in the time course analyses, due to fluctuations in access resistance occurring during ethanol application.

## Discussion

Here, using OVX female mice treated with E2 or VEH, we aimed to examine the role of E2 in modulating the response of VTA dopamine neurons to ethanol. In agreement with work by Vandegrift et al. [20], we found that E2 treatment of OVX mice potentiated ethanol excitation of VTA dopamine neurons without impacting the baseline firing rate of these cells. However, when examining synaptic inhibition onto the VTA dopamine neurons, as measured by spontaneous GABA_A_ receptor-mediated IPSCs, we did not observe any differences between treatment groups at baseline or as ethanol was applied. In fact, we observed substantial ethanol-stimulated synaptic inhibition (increased IPSC frequency) in the VTA dopamine neurons of both groups of OVX female mice, as others have shown for male mice [14,32,33]. We also observed a small effect of ethanol to increase IPSC amplitude, although again there was no difference between treatment groups. These findings do not support our initial hypothesis that E2 may promote ethanol excitation of VTA dopamine neuron firing by suppressing ethanol-stimulated GABAergic inhibition of these neurons. Nevertheless, this work contributes to the understanding of how sex and sex hormones influence the effect of ethanol on VTA dopamine neurons, and while our results did not support our hypothesis, they provide guidance for future investigation.

Generally, it has been difficult to clearly elucidate the mechanisms by which ethanol acts to influence dopamine neurons. While it seems paradoxical that both increased inhibition and excitation are driven simultaneously in the VTA by ethanol, it also indicates the apparent involvement of multiple independent mechanisms that are not yet fully understood. Moreover, VTA dopamine neurons are a heterogeneous population. Different subpopulations may be defined according to a number of criteria, such as anatomical localization, projection target, neurochemical content, or electrophysiological properties (e.g., I_h_ magnitude) [46–48], and various defined subpopulations can have differential responses to abused substances [49]. For example, other labs have reported that medial VTA dopamine neurons are more [50] responsive to ethanol compared to the more commonly studied lateral or canonical dopamine neurons, or that posterior VTA dopamine neurons are the ‘hotspot’ for ethanol activation [32]. Thus it is possible that we might have observed different effects of E2 if we had examined a different subpopulation, or a broader, more mixed population. We focused here on the classical lateral VTA dopamine neurons [51,52] and restricted our search via anatomical localization [43,53] by selecting neurons from a constrained region of the VTA (that focused on the parabrachial pigmented nucleus and avoided the paranigral, parafasicular, and the midline nuclei). Additionally, although we used hemizygous *Pitx3*-eGFP transgenic mice for ease in identifying dopamine neurons in slice, we also measured and compared I_h_ amplitude between groups. Medial VTA dopamine neurons tend to have little to no I_h_, and there may also be differences in I_h_ magnitude based on projection target [12]. Furthermore, variability in VTA dopamine neuron responses to ethanol has been shown to correspond to variability in I_h_ [54]. In our study, the distributions of I_h_ amplitude were similar between E2 and VEH treated animals. This suggests that we sampled from similar populations of dopamine cells, and that E2 facilitation of ethanol excitation of these cells was not mediated by an effect on I_h_.

Although we took measures to sample from a defined population of dopamine neurons, these neurons still receive diverse inputs [55], including several sources of inhibition. There are major GABAergic inputs from the NAc, the lateral hypothalamus, the medial preoptic area, and the rostromedial tegmental nucleus (RMTg) or tail of the VTA [56–59], and there is a substantial population of GABAergic interneurons within the VTA as well [16,60,61]. It is possible that E2 leads to increased firing in response to ethanol by selectively modulating the inhibition of one afferent source, which could have been obscured in our recording configuration. Unfortunately, we were not able to use a design which allowed us to discriminate the source of inhibition; however, tools are available that could facilitate these experiments and several groups have successfully targeted specific GABAergic inputs to the VTA [60,62–64]. There is strong evidence to point to the RMTg as a critical mediator of VTA dopamine neuron responses to ethanol, based on its well-described involvement in aversive components of drug and alcohol consumption and addiction [65–67]. Furthermore, there is evidence that RMTg to VTA projections are susceptible to control by different messengers compared to local inputs and NAc inputs [60,68,69]. Moreover, Melis and colleagues [70], when comparing a strain of alcohol preferring rats to their non-preferring counterparts, found that the magnitude and duration of VTA inhibition induced by stimulation of the RMTg was diminished in alcohol-preferring animals. However, given the functional and anatomical complexity within the striatum, a major input to and target of the VTA, it is likely that inputs from particular sub-regions of the NAc may also be critically important in modulating the activity of VTA dopamine neurons [71–73]. Of course, several new or overlooked pathways have been illuminated over the past decade, and there are several other substantial neurotransmitter inputs to the region, including glutamate and acetylcholine from pontine nuclei, serotonin from the dorsal raphe [56,57], and others, all of which could be directly or indirectly contributing to the activity of VTA dopamine neurons, local GABA neurons, and inputs to the region [74].

One factor that was not addressed in our experiments is the involvement of other inhibitory receptors. Recent work has demonstrated roles of the GABA_B_ receptor in controlling subsets of VTA neurons. Stephan Lammel’s group [75] showed that even within sub-regions of the VTA, dopamine neurons expressed and/or were manipulated by GABA_B_ receptors depending on both their projection targets and inputs. A small population of lateral VTA neurons that project to the lateral NAc receive inputs from medial NAc neurons that use GABA_B_ receptors to affect VTA dopamine neuron activity, separately from the rest of the reciprocal GABA_A_ receptor regulation. In a similar vein, there is evidence for the presence of glycine receptors on subsets of GABAergic terminals in the VTA [60] which, coupled with ethanol’s effects on the glycinergic system [76], provide another possible avenue to affect the firing rate [77]. While we did not attempt to directly test the involvement of either GABA_B_ or glycine receptors, these could be valuable experiments in future studies, particularly when examining the role of specific inputs and their associated mechanisms of inhibitory signaling.

Although we have detailed above some ways in which our experiments may not necessarily exclude the involvement of GABA signaling to dopamine neurons, other recent work does point to a role of glutamatergic signaling in the expression of E2/ERα-mediated enhancement of ethanol excitation of dopamine neuron firing. Vandegrift et al. [23] found that mGluR1 antagonism reduced ethanol excitation of VTA dopamine neurons in brain slices from E2-treated, but not VEH-treated, OVX mice, and from mice in diestrus, but not estrus. Thus, in addition to ERα, mGluR1 receptors are required to observe E2-mediated/estrous-cycle dependent enhancement of ethanol stimulation of dopamine neurons. Although ERα receptors express elsewhere in the VTA besides dopamine neuron soma, including interneuron soma and subsets of terminals projecting into the region [21,22,78], Vandegrift also found that selective knockout of ERα in dopamine neurons eliminated the enhanced responsiveness of those neurons during diestrus. Therefore, ERα signaling in dopamine neurons themselves is required for the observed enhancements of ethanol excitation. One possible interpretation of these findings is that ERα-mGluR heteromers on dopamine neurons may mediate the enhanced sensitivity to ethanol excitation. Indeed, there is a growing body of research demonstrating that estradiol can activate group I mGluRs and regulate neuronal intracellular signaling by, for example, enhancing cAMP-response element binding (CREB) protein phosphorylation [79,80]. The precise mechanisms by which promotion of CREB phosphorylation would enhance the sensitivity of VTA dopamine neurons to excitation by ethanol remain to be determined. However, it is clear from work done in other brain regions that manipulating CREB activity can have important consequences for neuronal membrane excitability[81]. Furthermore, CREB phosphorylation in the VTA and ER-mGluR heteromer signaling elsewhere in the brain have been implicated in sensitivity to drugs of abuse [27,28,82]. It is also possible that E2 increases ethanol excitation of dopamine neurons by promoting glutamatergic transmission more generally. E2 has been shown to potentiate the amplitude of ionotropic glutamate receptor responses and to promote glutamate release in several regions of the central nervous system [83–85].

Finally, while the present work provides further support for the idea that E2 may promote drinking behavior in females by influencing dopamine neuron responses to ethanol, it remains to be determined what roles E2 and estrogen receptors play in male mice. Vandegrift et al. [23] found that the same intra-VTA knockout of ERα that limited female ethanol drinking did not affect drinking by male mice, but it is not known whether E2 or estrogen receptors modulate dopamine neuron responses to ethanol in males, or whether E2 promotes drinking in males. Estradiol has in fact been shown to induce potentiation of hippocampal glutamatergic synapses in both sexes, albeit with different molecular signaling requirements [83]. Oberlander and Woolley [86] found that although different ER subtypes were engaged in males compared to females, both sexes experienced similar effects of E2 to increase pre- and post-synaptic glutamatergic signaling in the hippocampus. Notwithstanding differences in gonadal synthesis and circulating levels in the blood when compared to females, males can locally synthesize E2 in the brain from testosterone via aromatase [87,88]. This raises the possibility that E2 could influence VTA dopamine neuron excitation by ethanol and promote drinking in male mice, but via an ERα-independent mechanism. Recently, more researchers are starting to use the four-core genotype mice to probe genetic and gonadal components independently [89], which may allow for a clearer understanding of how circulating hormone levels interact with structural and organizational differences to express the full complement of sex differences. The increased use of this model may also help to illuminate how E2 and ERs function differently in males and females and sometimes, as indicated above, use different mechanisms to achieve the same end result.

Disentangling the mechanisms that underlie mesolimbic responses to ethanol and understanding how they may work together to produce behavioral output is a major task of the alcohol field. Our data did not support our hypothesis regarding the underlying mechanisms that could explain the increased excitation by ethanol following E2 treatment of OVX females, but there are some important caveats that we have discussed. Regardless, by replicating the original finding of Vandegrift et al., our experiments provided additional evidence in support of the idea that E2 plays an important role in modulating the physiological response to ethanol in the VTA of females. Taken altogether, these results begin to unravel the actions by which E2 may influence drinking behavior and prescribe some high-priority future experiments to address the lingering questions.

## Author Contributions

PL, AL and RM conceived and designed experiments. PL performed the experiments. PL and RM analyzed the data and interpreted the results, with some assistance from AL. PL and RM wrote the paper, and AL contributed with commentary and revisions.

## Acknowledgements

The authors wish to thank Heather Aziz and Jarrod Roach for technical assistance, and Rueben Gonzales, Mark Brodie, and Bertha Vandergrift for discussions and consultation regarding the execution of these experiments and/or this manuscript. Richard Morrisett is acknowledged for his role in funding acquisition.

